# Effects of maternal exercise modes on infant cord blood proteome

**DOI:** 10.1101/2024.11.11.623083

**Authors:** Filip Jevtovic, Breanna Wisseman, Fahmida Jahan, Alex Claiborne, David N. Collier, James E. DeVente, Steven Mouro, Tonya Zeczycki, Laurie J. Goodyear, Linda E. May

**Affiliations:** Department of Kinesiology, East Carolina University, Greenville, NC, USA; Human Performance Laboratory, East Carolina University, Greenville, NC, USA; East Carolina Diabetes and Obesity Institute, East Carolina University, Greenville, NC, USA; Section on Integrative Physiology and Metabolism, Joslin Diabetes Center, Harvard Medical School, Boston, Massachusetts, USA; Department of Kinesiology, University of Rhode Island, Kingston, RI, USA; Departments of Pediatrics, Brody School of Medicine, East Carolina University, Greenville, NC, USA; Obstetrics and Gynecology, Brody School of Medicine, East Carolina University, Greenville, NC, USA; Biochemistry and Molecular Biology, Brody School of Medicine, East Carolina University, Greenville, NC, USA

**Keywords:** maternal exercise, pregnancy, infant, proteomics, cord blood

## Abstract

The aim of this study was to show the effects of different maternal exercise modes on infant cord blood proteome. We used infant cord blood from two randomized controlled trials where women with a wide range of BMI and free of pregnancy complications participated in controlled and supervised aerobic, resistance, or combination (aerobic+resistance) exercise from <16 weeks of gestation until delivery. Results of this study showed that infant cord blood proteome is altered in a maternal exercise mode specific manner. Additionally, results showed 61 downregulated proteins common to all exercise modes, which correspond to gas transport, cellular stress response, reactive oxygen species metabolism, and other biological processes. Collectively, these data demonstrate the differential effect of maternal exercise modes on infant cord blood proteome.

## Introduction

Developmental origins of health and disease (DOHaD) suggests that the maternal environment during gestation impacts fetal metabolic programing and subsequent health. Maternal exercise during pregnancy has many beneficial effects on infant health including improvements of infant body composition^1–6^, increase in energy expenditure^7^, mitochondrial functional capacity and insulin action^1,8,9^. Nonetheless, despite many recognized benefits of maternal exercise, data regarding how different exercise modes affect infant health remains limited.

Interest in the distinct impact of maternal exercise modes on infant health comes from 1) differential effects of aerobic, resistance, or combination training on cardiometabolic health in non-gravid populations, 2) substantially different metabolic demand and physiological responses to exercise modes, 3) recent evidence of mode specific adaptations of placental proteome (paper in review) and infant mesenchymal stem cell metabolism^1,8^. Furthermore, potential use of maternal exercise modes as precision medicine to optimize infant health improvements continues to be a topic of considerable interest.

Blood plasma proteins have a role in mediating physiological processes (e.g., response to exercise), and can be used as predictors of one’s health (e.g., association of specific proteins with disease outcomes)^10–15^. For example, circulating peptide leptin stimulates proopiomelanocortin (POMC) and inhibits AgRP neurons in the arcuate nucleus to regulate energy homeostasis, and has been readily associated with obesity ^16,17^. The assessment of circulating proteins in infant cord blood can provide an insight into underlying biological processes across tissues and inform us about the intrauterine development and metabolic status of the fetus. Here, we aimed to elucidate how different modes of maternal exercise during gestation affect infant cord blood proteome.

## Methods

### Ethics statement

This study used infant cord blood collected in the ENHANCED (Enhanced Neonatal Health and Neonatal Cardiac Effect Developmentally) and EMCOR (Pregnancy Exercise Mode Effect on Childhood Obesity) studies (ClinicalTrials.gov Identifiers: NCT03838146 and NCT04805502). Approval for this study and all experiments was obtained from the East Carolina University Institutional Review Board and informed consent was obtained from each participant upon enrollment.

### Pre-intervention testing

All methods have previously been described. In brief, healthy females between 18 and 40 years of age and <16 weeks’ gestation were recruited from local obstetric clinics via flyers. Inclusion criteria included having a singleton pregnancy; free from chronic conditions including diabetes, hypertension, HIV, and others; not taking medications that may affect fetal development (e.g., antibiotics, SSRI); free of tobacco products, alcohol, and recreational drug use. After receiving clearance from the participant’s obstetric provider, participants were randomly assigned (via sealed, sequentially numbered envelopes derived from computer-generated randomization, Graph-Pad Prism Software) into an aerobic exercise, resistance exercise, a combination exercise, and non-exercising control group.

Participants completed a submaximal modified Balke treadmill test following the previously described methods for pregnant participants^18^ to measure peak oxygen consumption (VO_2peak_), and to determine target heart rate (THR) zones for subsequent exercise training. THR zones corresponded to maternal HR at 60-80% of maximal oxygen consumption, reflecting moderate intensity ^18^. For participants recruited during the COVID-19 pandemic, to minimize exposure and potential risk associated with exercise testing, THR was determined based on the pre-pregnancy physical activity level and age, using published guidelines ^18^. 21 participants had their THR zones estimated based on previously established protocols^18^, due to the COVID-19 pandemic; 16 participants had their VO_2peak_ assessed (n=3 control, n=6 aerobic, n=3 combination, n=4 resistance, no significant difference between groups, p>0.05). Importantly, cord blood protein expression was similar between

### Exercise intervention

Participants trained according to American College of Obstetricians and Gynecologists guidelines for approximately 24 weeks, beginning at 16 weeks’ gestation, and continued until delivery (∼40 weeks’ gestation)^19^. Participants performed moderate intensity (60%-80% maximal oxygen consumption and 12-14 rated perceived exertion) aerobic, resistance, or a combination exercise. Each exercise training session included a 5-minute warmup, 50-minute exercise period, and a 5-minute cooldown. Every session was supervised. Additionally, we used HR monitors to ensure that participants in all exercise groups (AE, CE, RE) maintained their HR within their assigned THR zone during each exercise session. Resistance training was done using free weights and seated Cybex machines. Aerobic training was done using treadmills, ellipticals, recumbent bicycles, or stair-stepping equipment. CE group spent the first half of the session doing resistance training followed by aerobic training. The CTRL group performed supervised stretching, breathing, and flexibility exercises at low intensity (<40% VO2peak). Maternal exercise adherence was calculated by dividing the number of sessions attended by the total number of possible sessions within the participants’ gestational period. Maternal exercise intensity (METs) was based on the published compendium of physical activity for the exercise performed in each session ^20^. The average maternal exercise dose during each week, expressed as METmin/wk, was quantified (frequency X duration of session) and then multiplied by the intensity (METs) of their exercise. Further, the total volume of exercise during pregnancy (Total METmin) was calculated by multiplying the METmin/wk by the total number of weeks of gestational exercise. Importantly, average, and total exercise volumes were calculated between 16-36 weeks of gestation, to avoid the influence of different gestational lengths between mothers (37-41 weeks) on exercise volume. The ability of our protocol to elicit maternal physiological adaptations to exercise has been previously reported ^21^.

### Maternal and infant birth measurements

Maternal measurements were obtained as previously described^22^. Maternal age, parity, pre-pregnancy weight and height and body mass index (BMI, kg/m^2^), gestational diabetes mellitus status (yes or no), length of gestation, and mode of delivery were abstracted from various sources including pre-screening eligibility and postpartum questionnaires as well as maternal and neonatal electronic health records. At 16 weeks of gestation, we obtained maternal BMI and determined maternal percent body fat via validated skinfold technique and age-adjusted equations^23,24^. Additionally, maternal fingerstick blood was analyzed using Cholestech LDX Analyzer (Alere Inc., Waltham, MA, USA) and point of care Lactate Plus Analyzer (Nova Biomedical, Waltham, MA, USA) to quantify maternal lipids (total cholesterol (TC), triglycerides (TG), HDL, non-HDL, LDL), glucose and lactate. <u>Birth measurements</u> (weight, length, Ponderal Index, abdominal, head and chest circumferences, 1 and 5-minute Apgar scores) and infant sex were extracted from neonatal electronic health records.

### Cord blood collection and proteomics

Cord blood was collected during delivery in anticoagulant tube, centrifuged to separate plasma and red blood cells, and stored in -80°C until analysis. Undepleted proteins from plasma samples (5 µL) were subsequently precipitated using ice-cold methanol (3:1 v/v) at -20° C and pelleted by centrifugation. The pellets were washed with ice cold methanol (x2) and allowed to air dry before further use. PreOmics iST kits (PreOmics GmbH) were used to prepare mass spectrometry grade peptides; as such precipitated proteins were resuspended in the provided lysis/denaturing buffer and denatured and alkylated at 95 °C for 10 min in a 1.5 mL microcentrifuge tube. After cooling to room temperature, digest reagent was added, and proteins were digested for 2 hrs at 37° C while shaking (600 rpm). Samples were then loaded onto the provided microcolumns for purification. Peptides were eluted *via* centrifugation using the provided elution buffer, dried to completeness under an N2 stream, and resuspended in loading buffer (98:2 water:acetonitrile+0.1% formic acid) at a concentration of 0.25 mg/mL.

Peptides were analyzed by nanoLC-MS/MS using an UltiMate 3000 RSLCnano system coupled to a Q Exactive Plus Hybrid Orbitrap mass Spectrometer (ThermoFisher) via nanoelectrospray ionization. Peptides were separated using an effective linear gradient of 4-35% acetonitrile (0.1% formic acid) over 135 min. For data-dependent acquisition in positive mode, MS spectra were acquired in positive mode. MS1 was performed at a resolution of 70,000 with an AGC target of 2×10^5^ ions and a maximum injection time of 100 ms. MS2 spectra were collected on the top 20 most abundant precursor ions with a charge >1 using an isolation window of 1.5 m/z and fixed first mass of 140 m/z. The normalized collision energy for MS2 scans was 30. MS2 spectra were acquired at 17,500 resolutions with a maximum injection time of 60 ms, an AGC target of 1×10^5,^ and a dynamic exclusion of 30 sec.

FragPipe (v 19.1)^25,26^ was used for raw data analysis with default search parameters for open and Label-Free Quantification Matching Between Runs (LFQ-MBR) workflows. Cord blood serum was searched individually with biological replicates (n=10) for each exercise group identified. An initial open search against the canonical + isoforms Uniprot Homo sapiens reference proteome (UP000005640, accessed 11/2023) was used to identify potential post-translational modifications for inclusion in the LFQ-MBR workflow. Precursor m/z tolerance was set to -150 to 500 Da and fragment tolerance was ± 20 ppm with 3 missed cleavages for Tryp and Lys-C allowed. Peptide spectrum matches (PSMs) were validated using PeptideProphet and results were filtered at the ion, peptide, and protein level with a 1% false discovery rate (FDR). Based on these initial searches, the following variable modifications were included in the LFQ-MBR analysis: oxidation (+15.5995 Da on Met), deamidation (+0.98401 Da on Gln and Asn), and fixed modification carbamidomethyl (+57.025 Da on Cys). For LFQ-MBR analysis, data were searched against the canonical Uniprot Homo sapiens reference proteome (UP000005640, accessed 11/2023 and 1/2024). Precursor ion m/z tolerance was ± 20 ppm with 3 missed cleavages for Trypsin/LysC allowed. The search results were filtered by a 1% FDR at the ion, peptide, and protein levels. PSMs were validated using Percolator and label-free quantification was carried out using IonQuant^27^ Match between runs FDR rate at the ion level was set to 10% for the top 300 runs. Proteins with >95% probability of ID, >2 unique peptides, and in more than 80% of a sample group (i.e. 7/10 injections) were considered high-confidence IDs and retained for analysis. Intensities were log2 transformed and normalized to the median intensities of the sample group. Relative abundances for low sampling proteins were determined via normal distribution in Perseus^28^.

### Statistical analysis

Maternal and infant characteristics were compared using one-way ANOVA. In case of not normally distributed data (e.g., parity) Kruskal-Wallis test was performed and data was expressed as median (min, max). For comparison of protein abundance between specific exercise groups and the control group (e.g., aerobic vs control), we used multiple unpaired t-tests with Welch correction for individual variance for each group, with p-value <0.1 for significance. Furthermore, Log2 fold change parameters for ‘meaningful’ differences between groups were set apriori for protein expression greater than 1.2 or less than -0.8 fold change. Statistical analyses were performed using GraphPad Prism version 9.3 (GraphPad Software, San Diego, CA) for Windows.

## Results

### Participant characteristics

Maternal and infant characteristics are presented in Table 1. All exercise groups had higher average weekly (MET•min/wk) and total pregnancy exercise volume (MET•min) compared to the control group. All groups had similar pre-pregnancy BMI, body fat percentage, and blood lipids, glucose, and lactate levels at 16 weeks of gestation. All participants were free of gestational diabetes. There was a similar infant sex distribution across the groups and all infants had similar birth weight, anthropometrics, and APGAR scores.

**Table 1.**
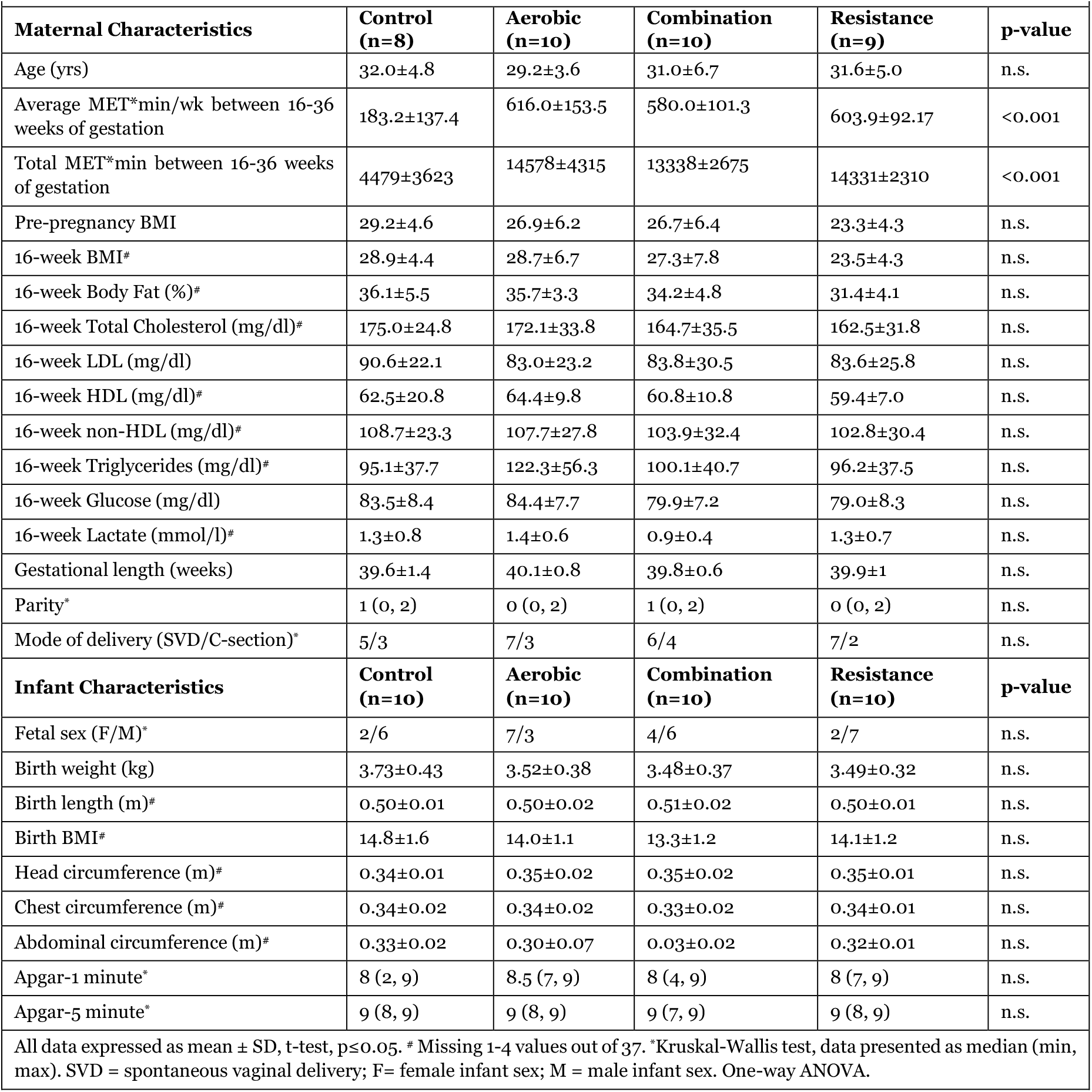
Maternal and Infant Characteristics.

### Effects of different maternal exercise modes on infant cord blood proteome

From the cord blood, we identified 324 proteins with high-confidence across all groups. For the aerobic, resistance, and combination groups, we observed 168, 115, and 108 proteins with statistically significant (p<0.1) differential abundance when compared to the control group, respectively. When significantly different proteins were restricted to a fold change greater than a 1.2 or less than -0.8 Log2 fold-change in abundance, we identified 159 proteins in aerobic, 94 in resistance, and 96 in combination group. Volcano plots of these proteins are presented in Figure 1A-C. Interestingly, there was only 1-2 significantly more abundant proteins in exercise, compared to the control group, while all other significantly different proteins were downregulated. Further, we used ShinyGO 8.0 (<u>ShinyGO 0.80 (sdstate.edu)</u>) to identify GO biological processes that these lower in abundance proteins correspond to (Figure 1D-F). Here, categories represent the annotation of enrichment of specific proteins, with fold-enrichment representing the magnitude of the pathway enrichment (number of genes altered / total number of genes in the pathway).

**Figure 1.**
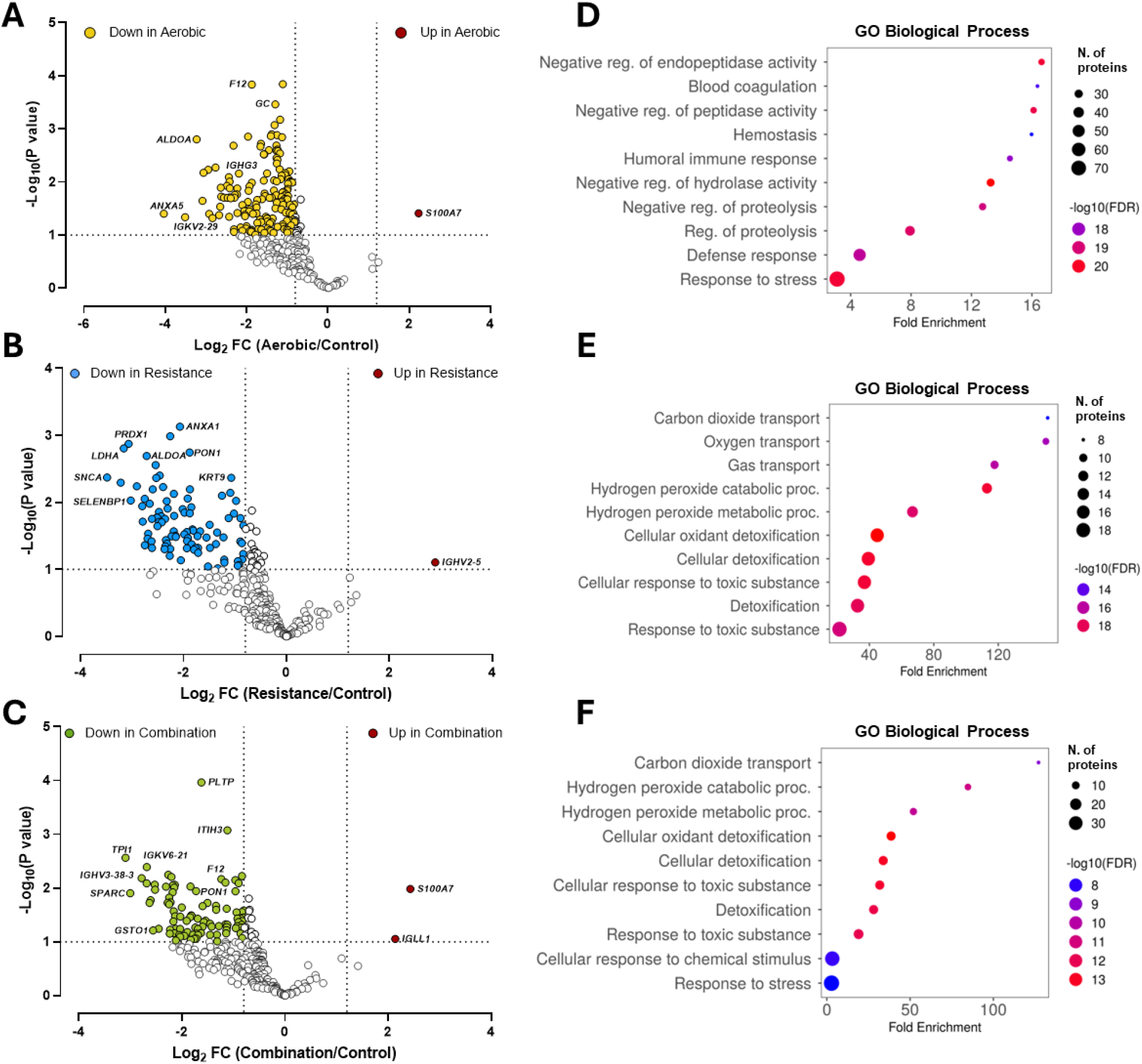
Effects of maternal exercise modes on the infant cord blood proteome. (A-C) Volcano plots of Log2 fold-change of protein abundances in exercise compared to the control groups vs. -Log10 (p-value) for the fold-change. Yellow, blue, and green symbols indicate proteins with significantly lower abundances (FC<-0.8) in the aerobic, resistance, and combination groups compared to control the group, respectively. White symbols show not significantly different proteins between exercise groups and the control group. (D-F) GO Biological Process enriched pathways corresponding to significantly less abundant proteins in exercise groups compared to the control group.

### Comparison of the effects of different maternal exercise modes on infant cord blood proteome

Comparing differentially abundant proteins identified as having significant and relevant changes in each exercise group relative to the control group, we identified 61 proteins common to all 3 exercise groups (Figure 2A). These 61 commonly downregulated proteins corresponded to GO Biological Process pathways involved in carbon dioxide and oxygen transport, hydrogen peroxide metabolic processes, cellular oxidant detoxification, reactive oxygen species metabolic processes,etc. (Figure 2B). The heatmap in Figure 2C shows the comparison of the exercise modes specific effect on these commonly altered proteins. Out of 61 proteins there were only 4 proteins where we observed exercise mode differences. Specifically, T-complex protein 1 subunit eta (CCT7, Q99832) and Immunoglobulin kappa variable 1-6 (IGKV1-6, A0A0C4DH72) were higher in combination compared to aerobic group (p<0.1). Additionally, triosephosphate isomerase (TPI1, P60174) was lower (p<0.1) in combination compared to aerobic group, and phospholipid transfer protein (PLTP, P55058) was higher in resistance compared to aerobic group (p<0.1). Collectively, these results show that despite differential effects of maternal exercise modes on infant cord blood proteome, there is still a significant portion of the proteome that is commonly altered across all exercise modes.

**Figure 2.**
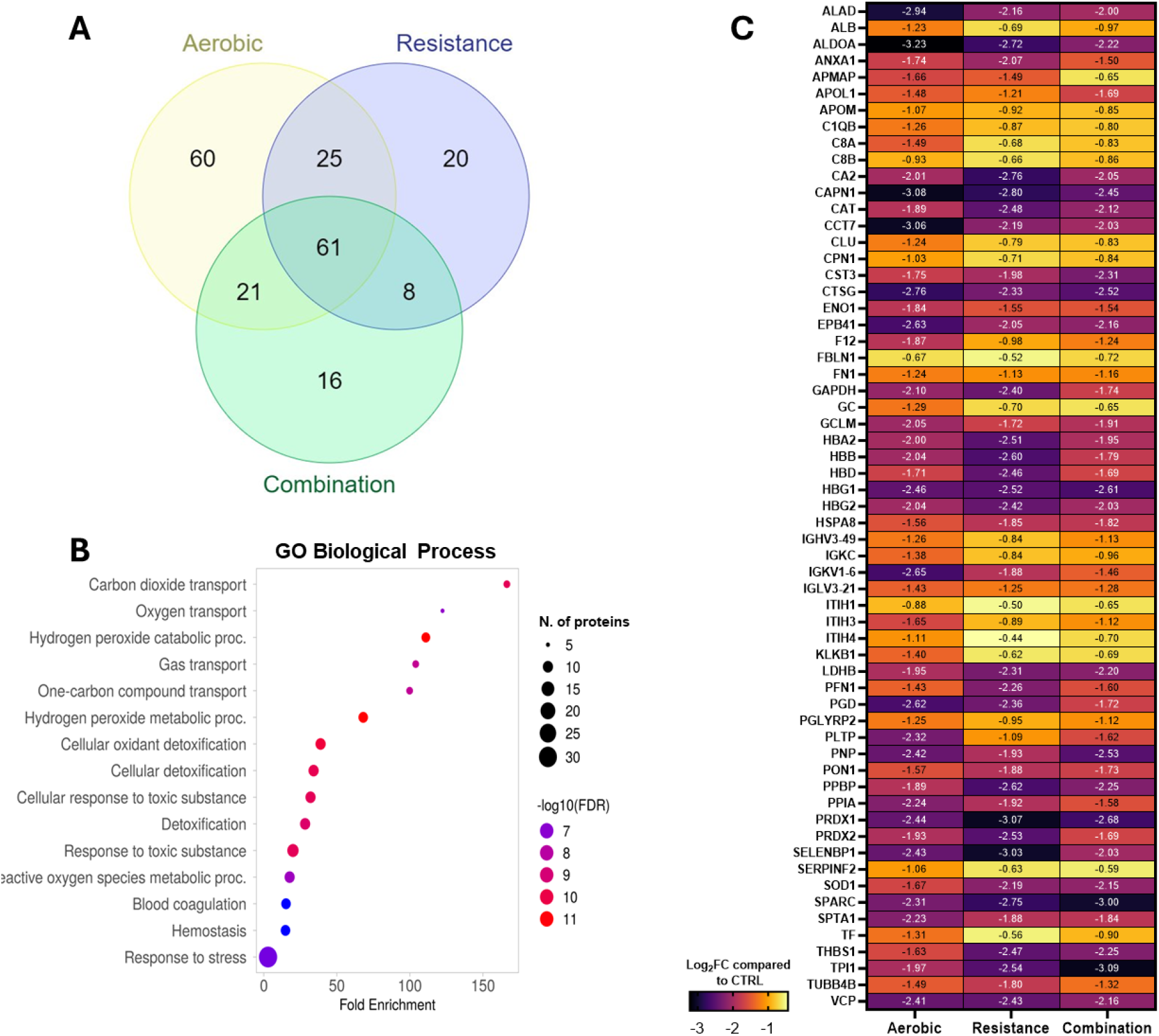
Commonly altered infant cord blood proteins across all maternal exercise modes. Venn diagram showing differentially altered proteins across exercise modes compared to the control group (A). GO Biological Process enrichment pathways that 61 commonly altered proteins across all exercise groups correspond to (B). Heatmap showing the exercise mode specific effect on 61 significantly altered proteins across all modes (C).

### Comparison between maternal exercise modes

To further examine these mode-specific dependencies, we compared the significant and relevant protein abundance differences arising from exercise group comparisons. Comparison of combination to aerobic exercise revealed 36 differentially abundant proteins (Figure 3A), with 9 upregulated and 4 downregulated proteins within Log^2^ fold-change below 0.8 and above 1.2 (Figure 3D). Further, there were 5 downregulated and 12 meaningfully upregulated proteins in resistance compared to aerobic control (Figure 3 B, E), and 19 lower in abundance and 6 proteins with higher abundances in combination compared to resistance exercise groups (Figure 3C, F). When comparing how these differences between groups overlap, we did not find any proteins common to all comparison (Figure 3G). Together these data demonstrate unique differences in protein abundance between maternal exercise modes.

**Figure 3.**
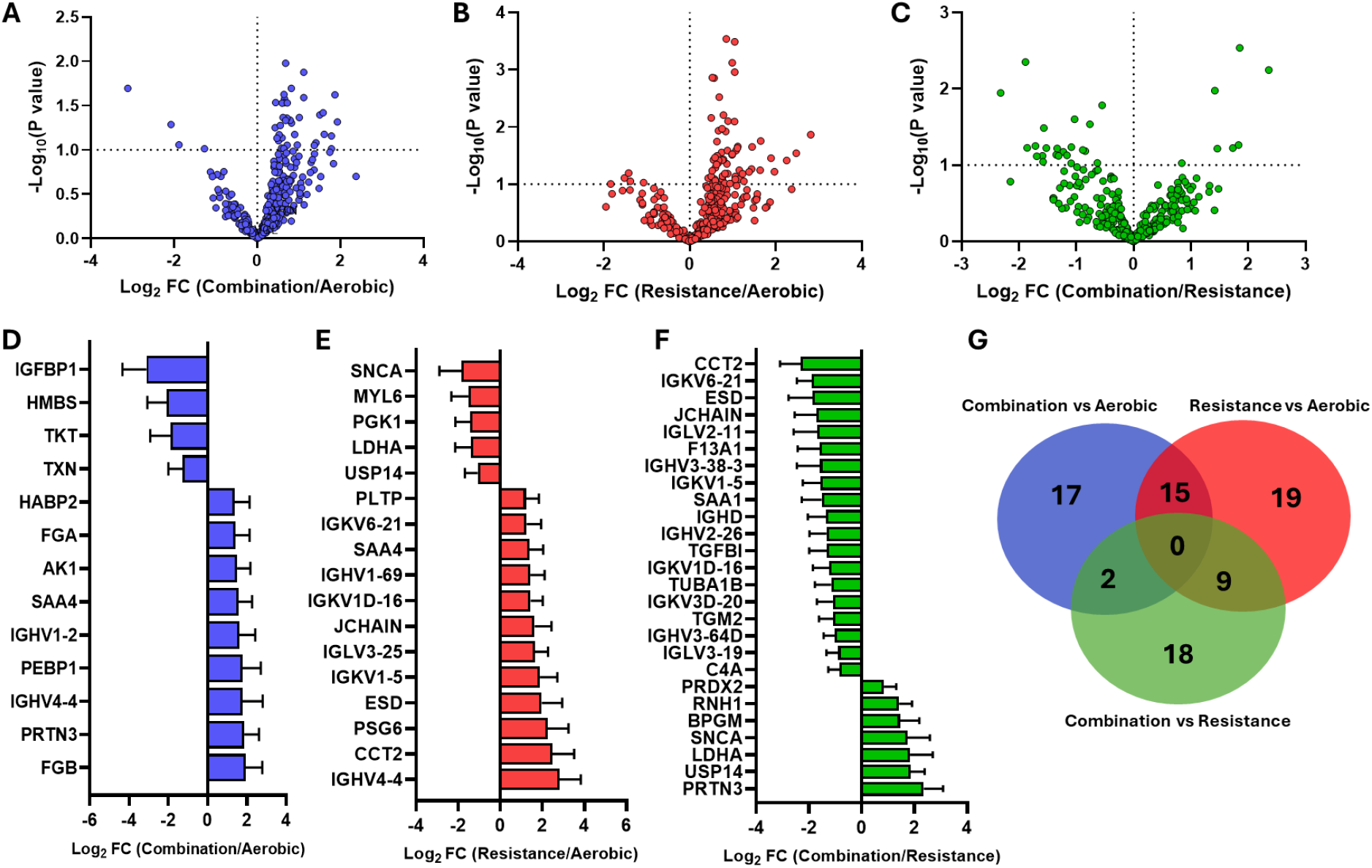
Vulcano plots with differentially expressed proteins between exercise groups (A-C) with significant proteins within the fold-change parameters shown below (D-F). Venn diagrams of identified proteins in different modes of exercise (G).

## Discussion

Understanding serum proteomic changes in response to maternal exercise provides a valuable insight into underlying infant physiological adaptations. Here we assessed how infant cord blood proteome changes in response to different modes of maternal exercise. Our results indicate that maternal exercise has a mode-specific effect on infant serum protein abundances, with only 61 commonly altered proteins across exercise modes when compared to the control group. Additionally, when comparing proteomic differences arising from exercise group comparisons, we did not observe any common proteins. Collectively, these data support the notion that maternal exercise mode is a significant factor that should be considered when prescribing maternal exercise, and that future research should aim to disentangle mechanisms behind differential physiological response to exercise modes.

While it is known that exercise is beneficial, it has recently been recognized that these benefits can vary depending on the type of exercise being performed. For instance, according to a preclinical study using a rat model of cardiac ischemia-repurfusion injury, different modes of exercise such as aerobic, resistance, and combination led to similarly reduction in infarct size ^29^, however, another study using a similar rat model of cardiac ischemia-repurfusion injury demonstrated that left ventricular performance was greater with high intensity aerobic exercise compared to resistance exercise^30^. In human studies, resistance exercise has been shown to better preserve muscle mass and quality in older adults with obesity^31^. Further, middle-aged women who performed both aerobic and resistance exercise improved their sympathetic nerve function; however, resistance exercise showed greater improvement in sympathetic and parasympathetic function suggesting resistance exercise is more beneficial in this population^32^. Further, gestational resistance exercise training was noted to be the most beneficial in improving systolic blood pressure, when compared to aerobic or combination training ^21^. Additionally, resistance exercise during pregnancy increased infant MSC glucose oxidation significantly more than aerobic exercise^1,8^. Taken together, the benefits of exercise modes differ, possibly due to the difference in the underlying mechanisms, and depending on the physiological context. Consistent with previous studies, here we showed that maternal exercise modes differentially modulate infant cord blood proteome. While it is difficult to interpret which mode of exercise is more beneficial for infant health, or how the differential effects of exercise modes on infant cord blood proteome will affect infant health later on, it is apparent that all exercise modalities during pregnancy improve maternal and offspring health^5,6^. Preclinical studies using rodent models could be useful in understanding the mechanisms of different modes of exercise on maternal and fetal tissues and how they affect health of the offspring in adulthood. In humans, long term follow-up studies in children from exercised mothers are needed to confirm which mode of exercise is more beneficial.

Free radicals, particularly reactive oxygen species, are produced during normal physiological processes and have a key role in intracellular signaling. Predominantly, reactive oxygen species such as superoxide and hydrogen peroxide, production is in mitochondria^33^. To maintain balance and minimize the deleterious effects of oxidative stress (e.g., on proteins, lipids, DNA), free radicals are readily scavenged by the enzymatic and non-enzymatic antioxidants. Further, production and antioxidant capacity are seemingly dependent on many factors including mitochondrial capacity, physical fitness, and even exercise performance ^34^. When it comes to exercise training, it has been well documented that chronic participation in exercise decreases blood and tissue oxidative stress, seemingly by increasing the efficiency of the antioxidant system(s)^35,36^. Here, across all exercise groups, we observed a decrease in the abundance of proteins corresponding to reactive oxygen species metabolism, detoxification, hydrogen peroxide catabolic processes. While we cannot comment on the activity of these antioxidants, it is interesting to observe a decrease in their protein abundance. Nonetheless, as we have previously observed significant improvements in infants MSC, a proxy of infants tissue, mitochondrial functional capacity1, it is possible that improvements of underlying infant metabolism could contribute to this effect; however, this necessitates further investigation.

The current findings highlight the susceptibility of the infant cord proteome to different maternal exercise modes. Strengths of this study are a prospective, randomized controlled trial study design which provides the strongest evidence for causality. We further acknowledge a few limitations. Our sample consisted of “apparently healthy” pregnant women; however, we had a range of BMI, healthy weight, overweight, and obese, which helps the generalizability of our findings. Further, we did not account for participant dietary habits, or paternal obesity which could be confounding factors in proteome adaptations. Future directions should address these limitations and focus on determining the effects of maternal exercise modes before and during pregnancy on maternal, and infant metabolic health.

In conclusion, this study aimed to assess how the infant cord blood proteome is remodeled in response to different maternal exercise modes. Here, we show that cord proteome is altered in a maternal exercise mode-specific way. Finally, these data showcase a myriad of altered pathways which incite further investigation to determine how these changes affect infant health.

## Acknowledgments

We thank Lindsey Rossa and Caitlyn Ollmann for assisting with specimen collection and subject recruitment. We thank the subjects for their participation.

